# Shivering, but not adipose tissue thermogenesis, increases as a function of mean skin temperature in cold-exposed men and women

**DOI:** 10.1101/2025.02.24.639578

**Authors:** Lauralyne Dumont, Gabriel Richard, Romain Espagnet, Frédérique Frisch, Mélanie Fortin, Arnaud Samson, Jonathan Bouchard, Réjean Fontaine, Etienne Croteau, Serge Phoenix, Stéphanie Dubreuil, Brigitte Guérin, Éric E. Turcotte, André C. Carpentier, Denis P. Blondin

## Abstract

Skin cooling results in the activation of heat generating mechanisms to counteract heat lost to the environment. Here, we aim to understand the extent that variations in cold-stimulated heat production may be driven by differences in the contribution of shivering and non-shivering thermogenesis (NST) and the interaction with biological sex. Using a novel mean skin temperature clamping technique in healthy men and women, our data shows that cold-stimulated heat production rises with increasing shivering and myocardial oxidative metabolism in a skin temperature-dependent fashion. Shivering and myocardial thermogenesis were also moderately associated. In contrast, adipose tissue NST did not increase in a linear manner to reductions in skin temperature. Men and women displayed similar thermoregulatory responses, although women presented more pronounced shivering through a greater recruitment of lower-body muscles and greater number of motor units recruited. Thus, shivering contributes proportionally to cold-induced thermogenesis whereas adipose tissue thermogenesis displays an all-or-none response.

## INTRODUCTION

Humans exposed to a cold environment rely primarily on behavioral strategies to defend their body temperature. When these strategies are insufficient, endogenous heat production must increase to counteract the heat lost to the environment. Studies that have simultaneously measured metabolic heat production and shivering have shown that there is tremendous variability in both the magnitude of cold-stimulated heat production^1^ and shivering intensity between individuals^2–5^, even for the same mean skin temperature^6^. This variability in response could be due to acclimatization/acclimation status^7–12^, or genetics^13–19^. It may be equally explained by methodological differences in cooling modality^6,20–22^, cooling duration^2,23,24^, and stimulation temperature^1,2,7,25,26^, or interindividual differences in biophysical properties, recruitment of thermogenic processes or biological sex. Here, by standardizing the cooling stimulus using a novel mean skin temperature clamping technique, we aim to understand the extent to which variations in cold-stimulated heat production may be driven by differences in the contribution of shivering and adipose tissue non-shivering thermogenesis (NST) and whether biological sex interacts with these outcomes.

Thermoregulatory cold defense responses are triggered by different combinations of peripheral and central thermosensory input^27^. Skin cooling provides the initial stimulus by activating cold-sensitive thermoreceptors in primary sensory nerve endings distributed in the skin^28,29^. The integration of these thermosensory inputs by the central nervous system triggers cold- defence effectors such as cutaneous vasoconstriction to limit heat loss and stimulates heat producing mechanisms that include a combination of shivering and non-shivering mechanisms such as brown adipose tissue (BAT) thermogenesis^20,30^, glycerolipid-fatty acid cycling^25,31^ and possibly other similar futile cycles previously identified in rodents^32–34^. To gain greater control over thermogenic responses, we recently developed a closed-loop temperature control system which allowed us to carefully control the skin cooling stimulus^6^. Despite this careful control, we still reported significant inter-individual variations in heat production and shivering intensity when the mean skin temperature of lean, healthy men and women was maintained at 31, 29 or 27°C ^6^. This led us to hypothesize that perhaps the interaction between shivering and non-shivering mechanisms of heat production is driving differences in whole-body thermogenesis. To date, few studies have examined shivering and non-shivering mechanisms simultaneously. Further, BAT thermogenesis, the most studied form of non-shivering thermogenesis in humans, has largely been investigated under a single cold stimulus^20,25,30,35^. Consequently, whether adipose tissue thermogenesis changes in response to different levels of skin cooling, thereby influencing its contribution to whole-body heat production, remains unknown. The primary objective of the present study was to examine the relationship between shivering and NST in adipose tissue (brown and white adipose tissue) in response to different levels of skin cooling, achieved by clamping the skin temperature at either 30°C or 26°C vs. room temperature (RT). We also investigated the effect of skin cooling on myocardial oxidative metabolism and whether biological sex impacts the interaction between these thermogenic mechanisms.

## RESULTS

### Skin cooling enhances whole-body thermogenesis, with the perceived cold thermal sensation varying based on the intensity of the cold

To standardize the cooling stimulus, a computer-controlled closed-loop temperature control system was used to decrease mean skin temperature at a target temperature of 30°C or 26°C during the acute cold exposure sessions (Figure 1). The energy expenditure increased significantly in response to the cold, increasing in a temperature-dependent manner (Figure 2A, 2B, 2C). Men presented with a significantly higher energy expenditure at RT and throughout the cold exposure (Figure 2B, 2C). Compared to RT, the change in energy expenditure averaged over the final 60 minutes of the cold exposure was significantly greater (*P* < 0.0001) at mean skin temperature of 26°C (107 kcal/h [95% CI: 89 to 126]) than 30°C (31 kcal/h [95% CI: 13 to 49]) (Figure 2D). There were no sex differences in the cold-induced increase in energy expenditure during the two acute cold exposures (Figure 2E). At RT, the absolute energy expenditure was significantly higher in men than women, but no difference was observed at 30°C and 26°C (Figure 2F). After adjusting absolute energy expenditure values for body mass, there were no sex differences at ambient temperature and between the two cold exposure intensities (P = 0.970; Figure 2G).

**Figure 1.**
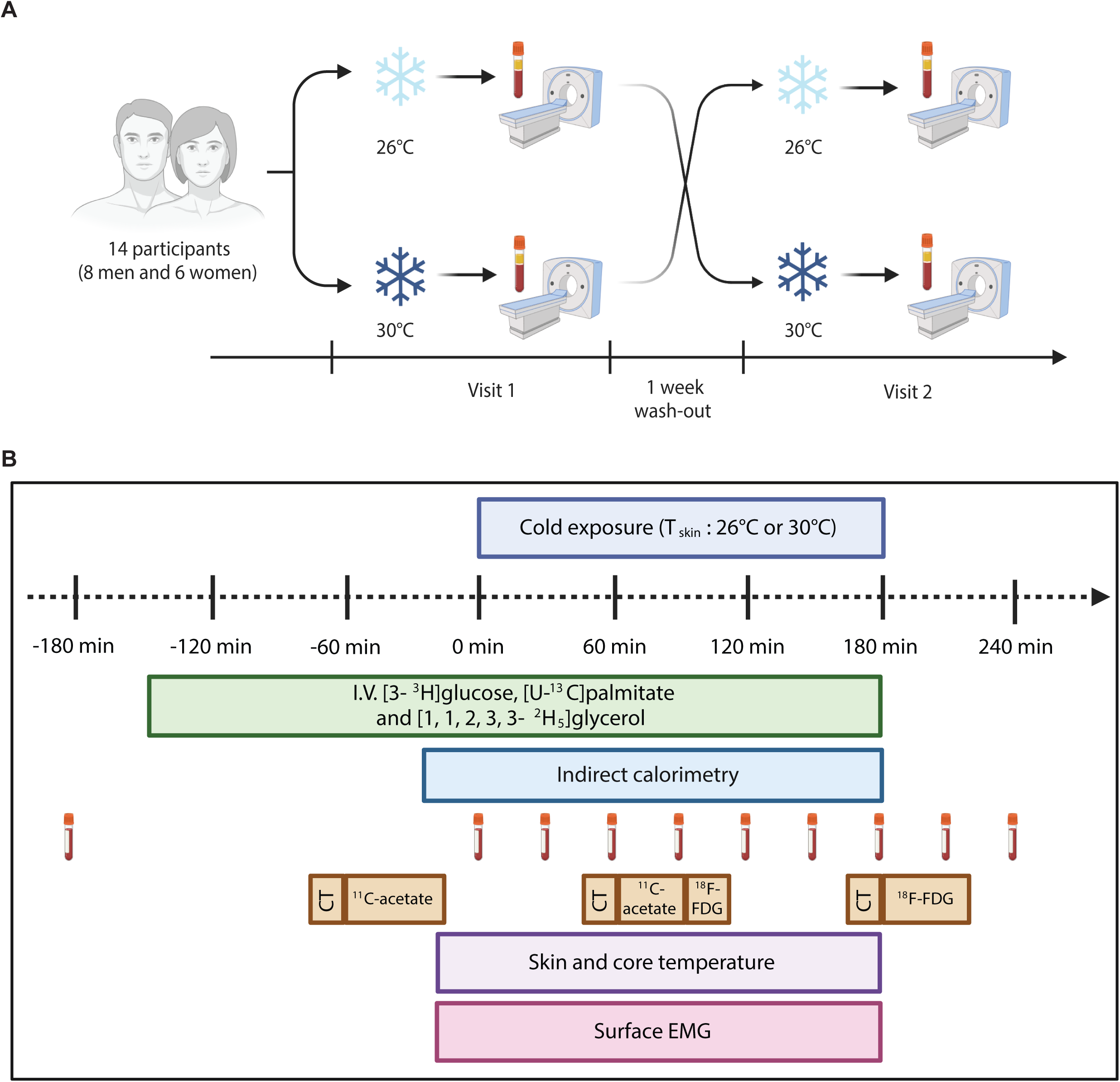
Overall study design. (A) Within-subject randomized crossover design (n = 14). Participants cooled to a target mean skin temperature of either 30°C or 26°C, with a 7-14-day wash-out period between conditions, using a custom-designed feedback control loop with a temperature and flow-controlled circulation bath. (B) Metabolic protocol for both study visits.

**Figure 2.**
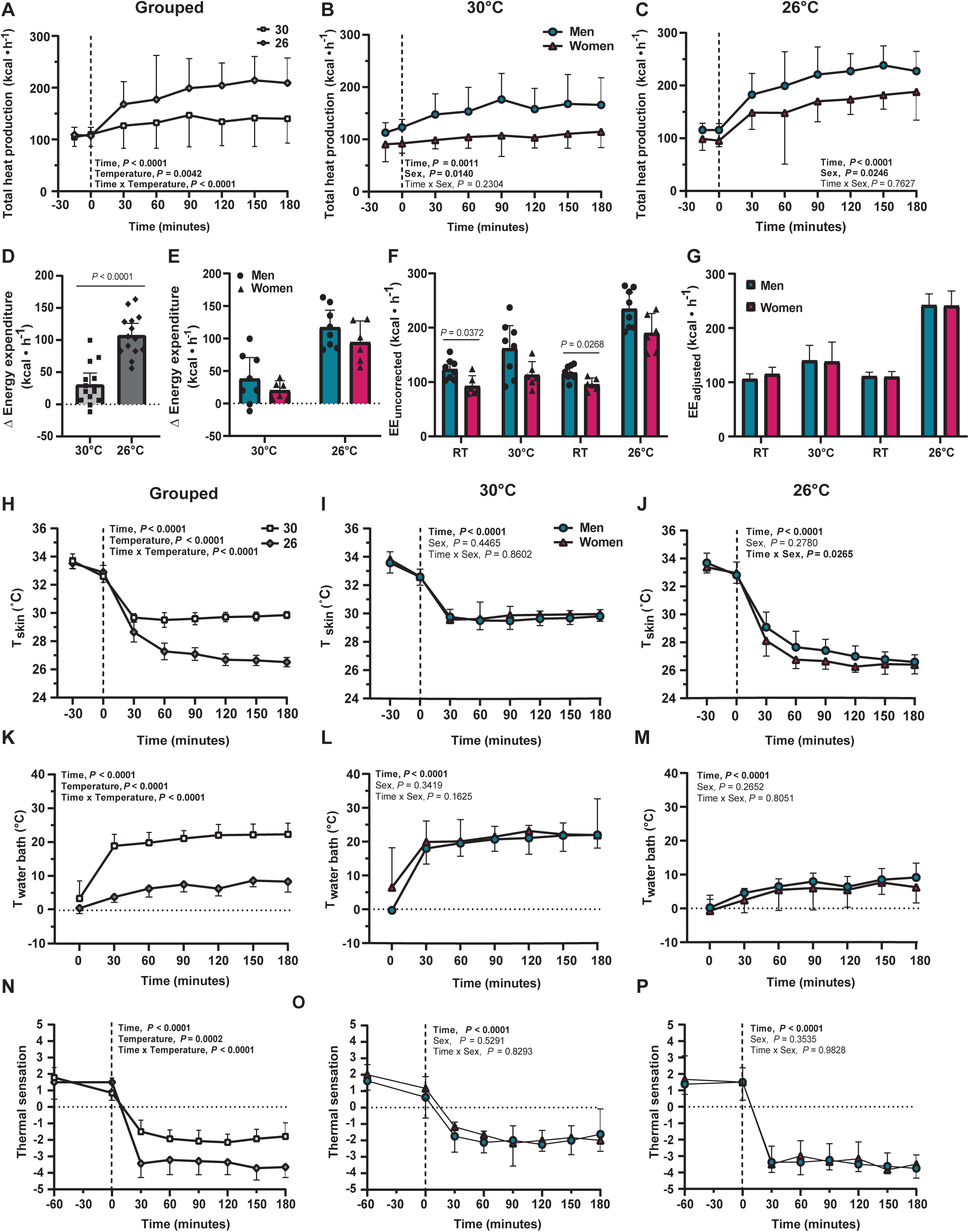
Skin cooling increases whole-body thermogenesis. (A-C) Whole-body energy expenditure over time among all participants (n = 14) (A), and in men (n = 8) and women (n = 6) when cooling to mean skin temperatures of either 30°C (B) or 26°C (C). (D and E) Cold-stimulated increase in energy expenditure when cooling to mean skin temperature of either 30°C or 26°C in all participants (n = 14) (D), and in men (n =8) and women (n = 6) (E). (F and G) Energy expenditure uncorrected (F) and adjusted for body mass (G) at room temperature and averaged over final 60 min of cold exposure when cooling to mean skin temperatures of either 30°C or 26°C in men (n = 8) and women (n = 6). (H-J), Mean skin temperature (Tskin), water bath temperature (Twater bath) (K-M), and thermal sensation over time (N-P) among all participants (30°C: n = 12-14, 26°C: n = 14) (left), and in men (n = 5-8) and women (n = 3-6) when cooling to mean skin temperatures of either 30°C (middle) or 26°C (right). Data are reported as means with 95% confidence interval (CI). P-values are shown only for significant differences (i.e. P < 0.05). A two-way ANOVA for repeated measures with Bonferroni post hoc test was used to determine the effects of the temperature, time and their interaction and a mixed-model ANOVA was used to include the between-subject effects of sex (A-C and E-P). The difference between 30°C and 26°C was determined using a paired-sample t-test (D). RT, room temperature; EE, energy expenditure; Tskin, skin temperature; Twater bath, water bath temperature.

The resulting changes in mean skin temperature and water bath temperature over time are presented in Figure 2H-M. With this protocol, mean skin temperature decreased quickly during the acute cold exposure, rapidly stabilizing to the target temperature of 30°C (29.8°C [95% CI: 29.5 to 30.1]) and 26°C (26.6°C [95% CI: 26.3 to 26.6]) among all the participants (*P* < 0.0001) with no change in the core temperature (Table 2). At mean skin temperature of 26°C, there was a transient time by sex interaction (*P* = 0.0265), resulting in a faster decrease in mean skin temperature for women compared to men (Figure 2J). The water temperature circulating through the suit increased progressively over time (*P* < 0.0001), stabilizing at a water temperature of 21.8°C (95% CI: 19.0 to 24.7) and 7.5°C (95% CI: 5.9 to 9.0) for the target mean skin temperature of 30°C and 26°C, respectively (Figure 2K). The water bath temperature increased over time for both men and women at a mean skin temperature of 30°C (*P* < 0.0001) and 26°C (*P* < 0.0001), with no statistically significant difference between the two (Figure 2L-M). There was a significant time by temperature interaction in thermal sensation (P < 0.0001) (Figure 2N). Compared with the room temperature period, participants reported feeling colder for both cold exposure temperatures. At a target skin temperature of 30°C, men and women reported feeling cold (*P* < 0.0001), while at 26°C, men and women reported feeling cold or very cold *(P* < 0.0001) (Figure 2O-P).

### Temperature-dependent increase in shivering intensity and selective muscle recruitment

Mean shivering intensity increased during the two cold exposure among grouped participants and in men and women (*P* < 0.0001) (Figure 3A-C). Mean shivering intensity averaged over the final 60 min of cold exposure increased with decreasing mean skin temperature (*P* < 0.0001) (Figure 3E) and increased to a greater extent in women than men, but only at a mean skin temperature of 26°C (temperature by sex interaction, *P* = 0.0102, Figure 3E). When controlling for differences in lean body mass, the mean skin temperature by sex interaction on shivering intensity was no longer statistically significant (*P* = 0.261).

**Figure 3.**
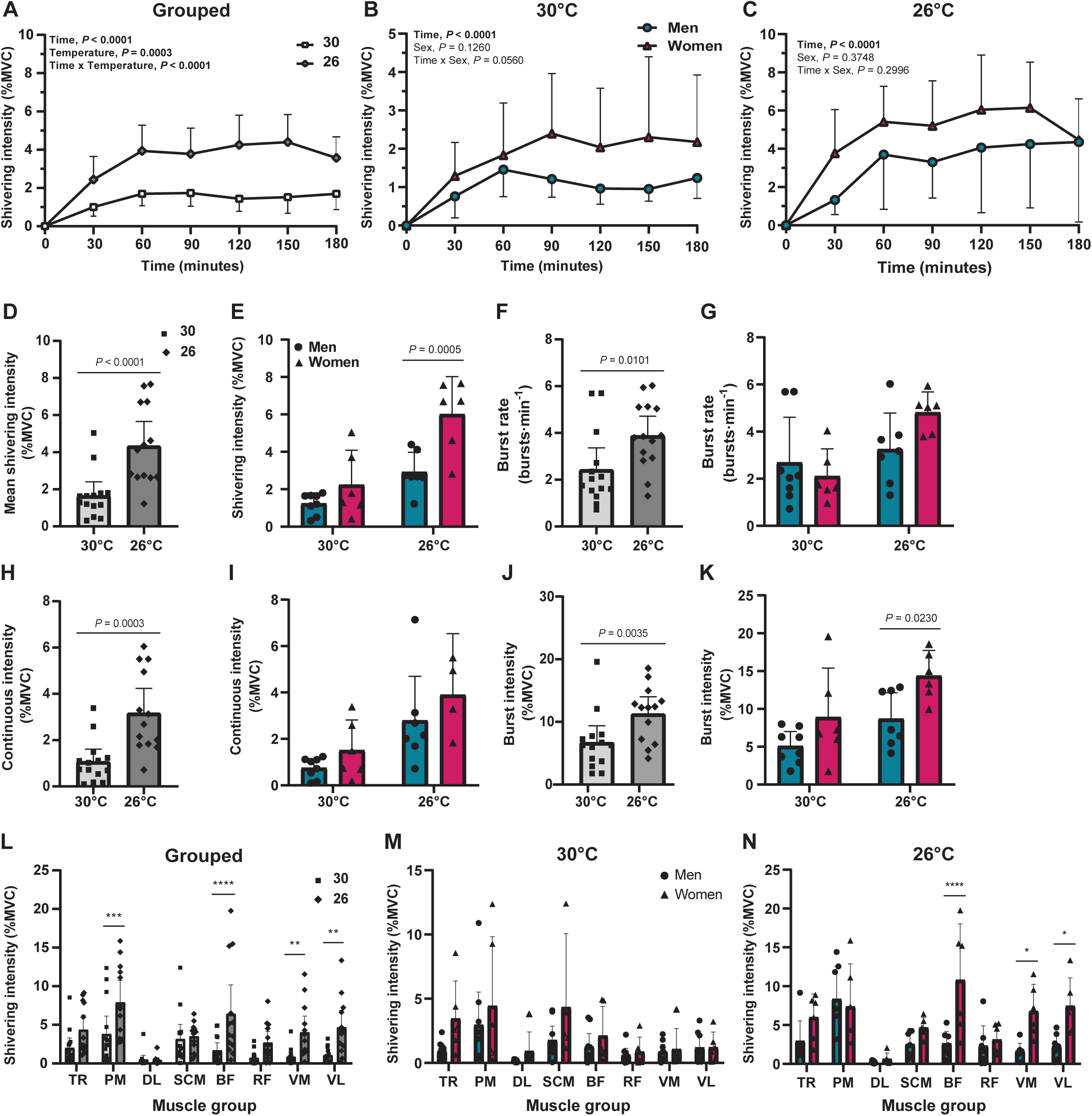
Temperature-dependent in shivering intensity and muscle recruitment. (A-C) Shivering intensity over time among all participants (30°C: n = 14, 26°C: n = 13) (A), and in men and women when cooling to mean skin temperature of either 30°C (men: n = 8, women: n = 6) (B) or 26°C (men: n = 7, women: n = 6) (C). (D-E) Mean shivering intensity averaged over final 60 min of cold exposure at skin temperature of either 30°C or 26°C among all participants (30°C: n = 14, 26°C: n = 13) (D), and in men (30°C: n = 8, 26°C: n = 7) and women (30°C: n = 6, 26°C: n = 6) (E). (F and G) Burst rate averaged over final 60 min of cold exposure at skin temperature of either 30°C or 26°C among all participants (30°C: n = 14, 26°C: n = 13) (F), and in men (30°C: n = 8, 26°C: n = 7) and women (30°C: n = 6, 26°C: n = 6) (G). (H and I) Continuous shivering intensity averaged over final 60 min of cold exposure at skin temperature of either 30°C or 26°C among all participants (30°C: n = 14, 26°C: n = 13) (H), and in men (30°C: n = 8, 26°C: n = 7) and women (30°C: n = 6, 26°C: n = 6) (I). (J and K) Burst shivering intensity averaged over final 60 min of cold exposure at skin temperature of either 30°C or 26°C among all participants (30°C: n = 14, 26°C: n = 13) (J), and in men (30°C: n = 8, 26°C: n = 7) and women (30°C: n = 6, 26°C: n = 6) (K). (L-N) Mean shivering intensity averaged over final 60 min of cold exposure in 8 different muscles [m. trapezius (TR), m. pectoralis major (PM), m. deltoideus (DL), m. sternocleidomastoid (SCM), m. biceps femoris (BF), m. rectus femoris (RF), m. vastus medialis (VM), and m. vastus lateralis (VL)] among all participants (30°C: n = 14, 26°C: n = 13) (L), and in men (30°C: n = 8, 26°C: n = 7) and women (30°C: n = 6, 26°C: n = 6) during mild cold exposure at skin temperature of either 30°C (M) or 26°C (N). Data are reported as means with 95% confidence interval (CI). P-values are shown only for significant differences (i.e. P < 0.05). A two-way ANOVA for repeated measures with Bonferroni post hoc test was used to determine the effects of the temperature, time and their interaction and a mixed-model ANOVA was used to include the between-subject effects of sex. The difference between 30°C and 26°C was determined using a paired-sample t-test (D, F, H, and J). MVC, maximal voluntary contraction; RT, room temperature.

The shivering pattern (continuous vs. burst shivering) of eight large muscles was continuously measured throughout cold exposure using surface electromyography (sEMG). In the final 60 min of cold exposure, the burst shivering rate (30°C: 2.4 bursts·min^-1^ [95% CI: 1.5 to 3.4] vs. 26°C: 4.0 bursts·min^-1^ [95% CI: 3.1 to 4.9]; *P* = 0.0101), continuous shivering intensity (30°C: 1.1% maximal voluntary contraction (MVC) [95% CI: 0.5 to 1.6] vs. 26°C: 3.2% MVC [95% CI: 2.1 to 4.2]; *P* = 0.0003) and burst shivering intensity (30°C: 6.7% MVC [95% CI: 4.1 to 9.4] vs. 26°C: 11.3 % MVC [95% CI: 8.6 to 14.0]; *P* = 0.0035) was lower at 30°C compared with 26°C (Figure 3F, H and J). There was no sex difference in these three parameters except for the burst shivering intensity during the coldest acute cold exposure (*P* = 0.0230), where the burst shivering intensity represented 8.7% MVC (95% CI: 5.3 to 12.1) in men and 14.4% MVC (95% CI: 11.0 to 17.7) in women (Figure 3G, I and K). Shivering intensity in *m. pectoralis major*, *m. biceps femoris*, *m. vastus medialis,* and *m. vastus lateralis* was significantly higher at a mean skin temperature of 26°C than mean skin temperature of 30°C (Figure 3L). There were no sex differences in muscle recruitment when the mean skin temperature was clamped to 30°C (Figure 3M). However, at a colder mean skin temperature, the shivering intensity was approximately 4-fold higher in women than in men for three lower body muscles: *m. biceps femoris* (women: 10.8% MVC [95% CI: 3.6 to 18.0] vs. men: 2.6% MVC [95% CI: 1.1 to 4.2]; *P* < 0.0001), *m. vastus medialis* (women: 6.8% MVC [95% CI: 3.3 to 10.2] vs. men: 1.7 % MVC [95% CI: 0.7 to 2.7]; *P* = 0.0297) and *m. vastus lateralis* (women: 7.5% MVC [95% CI: 3.8 to 11.1] vs. men: 2.3% MVC [95% CI: 0.8 to 3.8]; *P* = 0.0255) (Figure 3N).

### Myocardial oxidative metabolism increases with decreasing mean skin temperature

Compared to room temperature conditions, the change in diastolic and systolic blood pressure were similar between the two mean skin temperatures (Figure 4A-D). In the present study, while the heart rate decreased by 4 beats·min^-1^ (95% CI: -10 to 2) below room temperature levels when skin temperature was clamped at 30°C, it increased by 3 beats·min^-1^ (95% CI: -3 to 8) compared to room temperature levels at a mean skin temperature of 26°C, which was significantly different between the two temperatures (*P* = 0.0120). We determined the rate pressure product (RPP) to investigate how two skin cooling temperatures affect myocardial metabolic demands. Compared to room temperature, the change in RPP was greater when clamping mean skin temperature at 26°C than when clamping mean skin temperature at 30°C (1227 mmHg· beats·min^-1^ [95% CI: 277 to 2177] vs. 180.2 mmHg· beats·min^-1^ [95% CI: -430 to 790], *P* = 0.0099) (Figure 6G). The change in diastolic and systolic blood pressure, heart rate and RPP did not differ between sexes at mean skin temperature of either 30°C or 26°C (Figure 4B, D, F and H).

**Figure 4.**
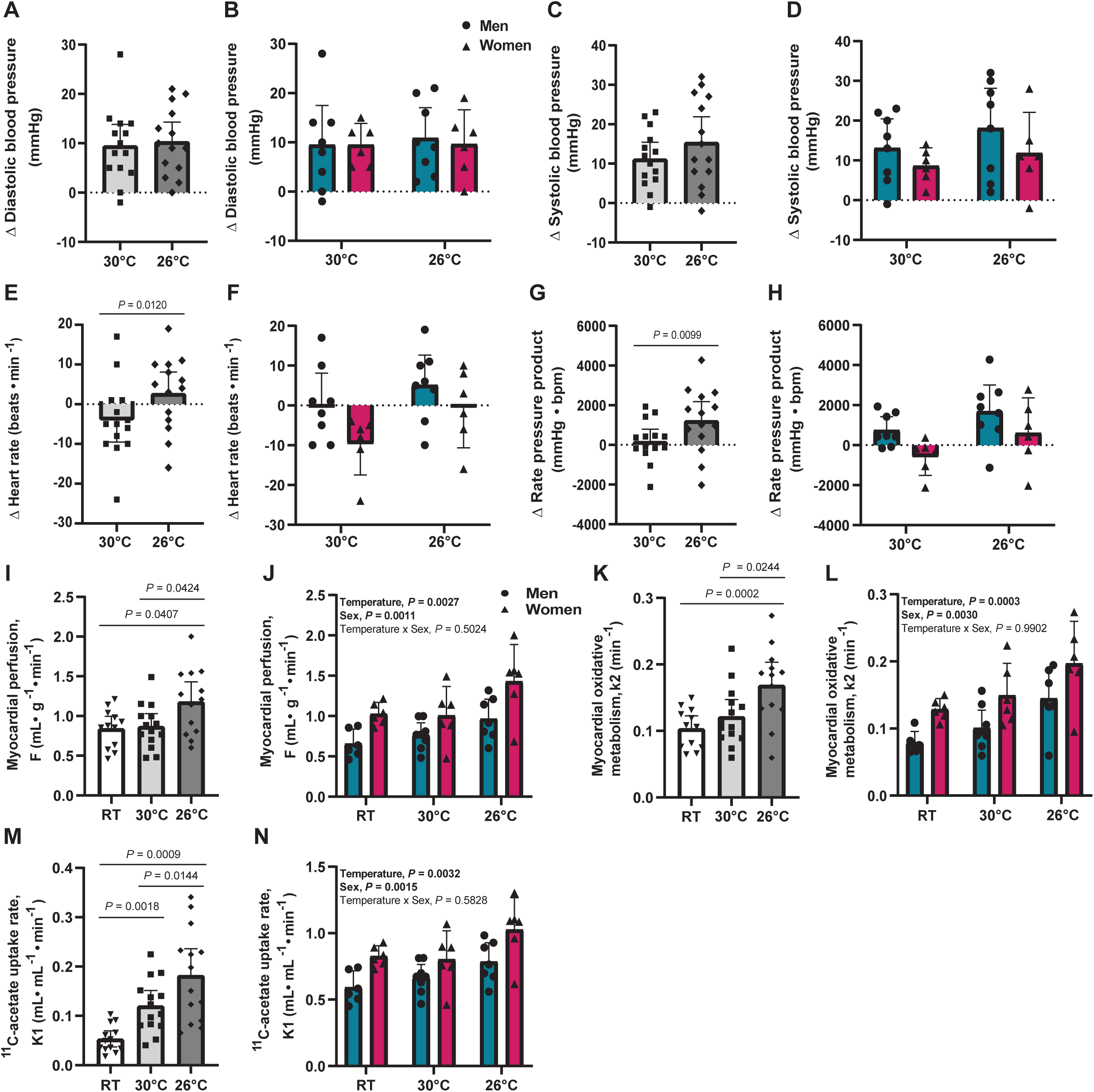
Myocardial oxidative metabolism increases as well as rate pressure product at the coldest mean skin temperature of 26°C. (A-H) Change in diastolic blood pressure (A and B), systolic blood pressure (C and D), heart rate (E and F), and rate pressure product (G and H) from room temperature levels in response to a mild cold exposure at skin temperature of either 30°C (n = 14) or 26°C (n = 14) among all participants (A, C, E and G), and in men (n = 8) and women (n = 6) (B, D, F and H). (I and J) Myocardial perfusion (F) obtained from the [11C]acetate analysis of the heart at room temperature (n =12) and in response to cold exposure at mean skin temperature of either 30°C (n = 14) and 26°C (n =14) among all participants (I), and in men (RT: n = 6, 30°C: n = 7, 26°C: n = 7) and women (RT: n = 6, 30°C: n = 6, 26°C: n = 6) (J). (K and L) Myocardial oxidative metabolism (k2) obtained from the [11C]acetate analysis of the heart at room temperature (n = 12) and in response to cold exposure at mean skin temperature of either 30°C (n = 14) and 26°C (n = 14) among all participants (K), and in men (RT: n = 6, 30°C: n = 7, 26°C: n = 7) and women (RT: n = 6, 30°C: n = 6, 26°C: n = 6) (L). (M and N)[11C]acetate uptake (K1) of the heart at room temperature and in response to cold exposure at skin temperature of either 30°C (n = 14) or 26°C (n = 14) among all participants (M), and in men (RT: n = 6, 30°C: n = 7, 26°C: n = 7) and women (RT: n = 6, 30°C: n = 6, 26°C: n = 6) (N). Data are reported as means with 95% confidence interval (CI). P-values are shown only for significant differences (i.e. P < 0.05). The difference between 30°C and 26°C was determined using a paired-sample t-test (A, C, E, and G). A one-way ANOVA for repeated measures with Bonferroni post hoc test was used to determine statistically significant differences between the three conditions (room temperature, 30°C, and 26°C) (I, K, and M). A mixed-model ANOVA with Bonferroni post hoc test was used to determine the effects of the temperature, the sex, and their interaction. RT, room temperature; bpm, beats per minute.

With the large PET scanner field of view (26 cm), this study was the first to simultaneously quantify myocardial and BAT oxidative metabolism in humans. Myocardial perfusion (F), oxidative metabolism (*k*_2_) and [^11^C]-acetate uptake rate (*K*_1_) were assessed by [^11^C]-acetate PET imaging. To do this, we analyzed PET dynamic images of the heart by drawing a ROI in the left ventricle with PMOD modeling tool, PMOD Kinetic Modeling (PKIN) using one compartment model (Card acetate, 1 compartment - PMOD Technologies, version 3.7). The myocardial perfusion (F) was significantly higher at a mean skin temperature of 26°C (1.181 mL·g^-1^·min^-1^ [95% CI: 0.931 to 1.431]) than at room temperature (0.842 mL·g^-1^·min^-1^ [95% CI: 0.688 to 0.996]; *P* = 0.0407) or at 30°C (0.870 mL·g^-1^·min^-1^ [95% CI: 0.709 to 1.030]; *P* = 0.0424) (Figure 4I).

The same results were observed for the myocardial oxidative metabolism (*k*_2_) (Figure 4K). Compared to room temperature (0.104 min^-1^ [95% CI: 0.085 to 0.123]; *P* = 0.0002) and mean skin temperature of 30°C (0122 min^-1^ [95% CI: 0.096 to 0.147]; *P* = 0.0244), the myocardial oxidative metabolism was greater when the mean skin temperature was clamped to 26°C (0.169 min^-1^ [95% CI: 0.134 to 0.203]). The [^11^C]-acetate uptake rate was significantly higher at mean skin temperature of 26°C (0.897 mL·mL^-1^·min^-1^ [95% CI: 0.763 to 1.031]) than 30°C (0.725 mL·mL^-1^·min^-1^ [95% CI: 0.627 to 0.822]; *P* = 0.0144), and was significantly higher at 26°C (*P* = 0.0009) and 30°C (*P* = 0.0018) than at room temperature (0.709 mL·mL^-1^·min^-1^ [95% CI: 0.611 to 0.203]) (Figure 4M). Myocardial perfusion (*P* = 0.0011), oxidative metabolism (*P* = 0.0030) and [^11^C]- acetate uptake rate (*P* = 0.0015) were significantly higher in women than in men at room temperature, and at mean skin temperatures of 30°C and 26°C (Figure 4J, L and N).

### Cold exposure increases BAT oxidative metabolism but not in a temperature-dependent fashion

Figure 4 presents the rates of BAT oxidative metabolism and tissue-specific glucose uptake, which were assessed using a multi-tracer approach with [^11^C]-acetate and ^18^F- fluorodeoxyglucose (FDG), respectively. [^11^C]-acetate uptake (*K*_1_, mL·g^−1^·min^−1^), a marker of tissue perfusion, oxidative metabolism (*k*_2_, min^−1^) and lipid synthesis (*k*_3_, min^−1^) were estimated using a four-compartment, two-tissue model previously developed specifically to estimate BAT kinetics ^36^. The rate of BAT [^11^C]-acetate uptake increased in the cold in a dose-dependent manner, being significantly higher at a clamped skin temperature of 30°C (0.121 mL·g^−1^·min^−1^ [95% CI: 0.089 to 0.152], *P* = 0.0010) and 26°C (0.182 mL·g^−1^·min^−1^ [95% CI: 0.128 to 0.236], *P* = 0.0005) compared to room temperature (0.054 mL·g^−1^·min^−1^ [95% CI: 0.038 to 0.069]). BAT [^11^C]-acetate uptake was also significantly different between the two cold exposure (*P* = 0.0144) (Figure 5A). BAT oxidative metabolism increased from 0.795 min^-1^ (95% CI: 0.519 to 1.070) at room temperature to 1.806 min^-1^ (95% CI: 1.361 to 2.250; *P* = 0.0019) and 1.909 min^-1^ (95% CI: 1.458 to 2.361; *P* = 0.00133), at mean skin temperature of 30°C and 26°C, respectively. BAT oxidative metabolism was not statistically different between 30° and 26°C (Figure 4B, *P =*0.8579). We found that at room temperature, BAT intracellular lipid synthesis was significantly greater than at mean skin temperature of 30°C (*P* = 0.0422) and 26°C (*P* = 0.0037) but did not differ between the two skin cooling conditions (Figure 5C). However, there was no difference in the change in CT- derived tissue radiodensity during cold exposure, a marker of intracellular triglyceride content (Figure 5G-I). Decreasing the mean skin temperature to either 30°C or 26°C did not elicit any sex- dependent differences in BAT [^11^C]-acetate uptake, oxidative metabolism, lipid synthesis or radiodensity (Figure 5D-F and H-I). The Patlak linearization model was used to evaluate the rate of net FDG uptake per gram of tissue in BAT, cervical subcutaneous WAT and skeletal muscles between the two mean skin temperatures (Figure 5J and L) in men and women (Figure 5K and M). During cold exposure, the fractional FDG uptake (*Ki*) in supraclavicular BAT, skeletal muscles and subcutaneous WAT was not different between the two clamped temperature (Figure 5J) and between men and women (Figure 5K). The rate of net FDG uptake in BAT at a mean skin temperature of 30°C and 26°C respectively increased to 58 nmol·g^-1^·min^-1^ (95% CI: 33 to 83) and 60 nmol·g^-1^·min^-1^ (95% CI: 33 to 88) (Figure 5L), which was greater than previously reported under room temperature conditions [9 ± 4 nmol·g^-1^·min^-1^, N=27]^37^. Mass-specific net FDG uptake was greater in BAT than skeletal muscles (*P* < 0.0001) and scWAT (*P* < 0.0001) for both mean skin temperature. We did not observe any sex differences in net FDG uptake in BAT, skeletal muscles and scWAT when the skin temperature was clamped to 30°C or 26°C (Figure 5M).

**Figure 5.**
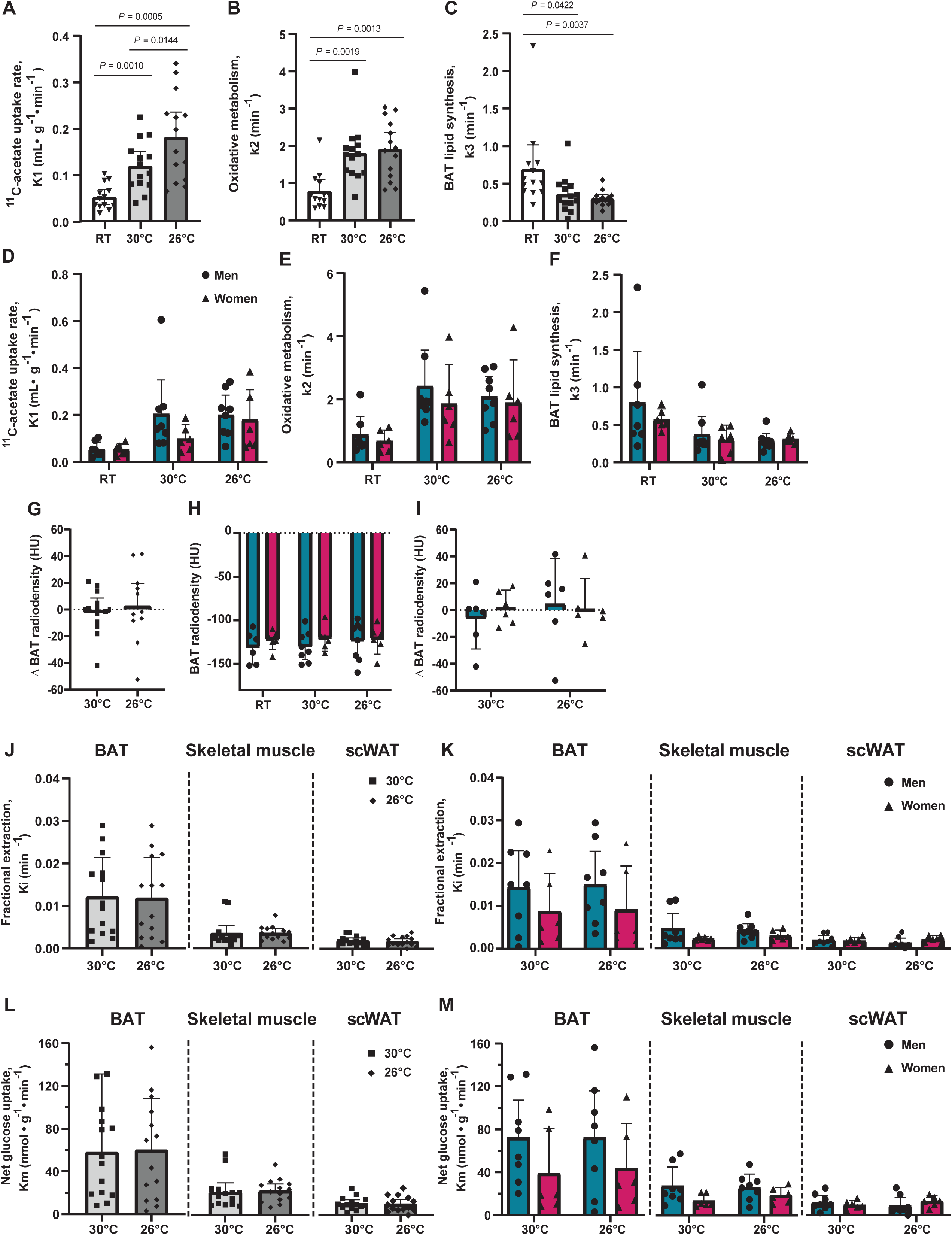
BAT oxidative metabolism increases in the cold, but not in a temperature- dependent fashion. (A-F) [11C]acetate uptake rate (A and D, K1), oxidative metabolism (B and E, k2), and lipid synthesis (D and F, k3) of supraclavicular BAT at room temperature (n = 13) and during mild cold exposure at skin temperature of either 30°C (n = 14) or 26°C (n = 14) among all participants (A-C), and in men (RT: n = 7, 30°C: n = 8, 26°C: n = 8) and women (RT: n = 6, 30°C: n = 6, 26°C: n = 6) (D-F). (G) Cold-stimulated change in supraclavicular BAT radiodensity at skin temperature of either 30°C (n = 12) or 26°C (n = 12) among all participants. (H) Supraclavicular BAT radiodensity at room temperature (men: n = 6, women: n =6) and in response to cold exposure at skin temperature of either 30°C (men: n = 8, women: n =6) or 26°C (men: n = 8, women: n =6) in men and women. (I) Cold-stimulated change in supraclavicular BAT radiodensity at skin temperature of either 30°C or 26°C in men (n = 6) and women (n = 6) (I). (J and K) Supraclavicular BAT, WAT, and skeletal muscle fractional glucose uptake in response to a mild cold exposure at skin temperature of either 30°C (n = 14) or 26°C (n = 14) among all participants (J), and in men (n = 8) and women (n = 6) (K). Skeletal muscle fractional glucose uptake represents the mean uptake of regions of interest drawn over the m. pectoralis major, m. trapezius, m. deltoideus, m. sternocleidomastoid, m. levator scapulae, m. latissimus dorsi, and m. erector spinae. (L and M) Supraclavicular BAT, WAT, and skeletal muscle net glucose uptake in response to a mild cold exposure at skin temperature of either 30°C (n = 14) or 26°C (n = 14) among all participants (L), and in men (n = 8) and women (n = 6) (M). Skeletal muscle glucose uptake represents the mean uptake of regions of interest drawn over the m. pectoralis major, m. trapezius, m. deltoideus, m. sternocleidomastoid, m. levator scapulae, m. latissimus dorsi, and m. erector spinae. Data are reported as means with 95% confidence interval (CI). P-values are shown only for significant differences (i.e. P < 0.05). The difference between 30°C and 26°C was determined using a paired-sample t-test (G, J, and L). A one-way ANOVA for repeated measures with Bonferroni post hoc test was used to determine differences between the three conditions (room temperature, 30°C, and 26°C) (A-C). A mixed-model ANOVA with Bonferroni post hoc test was used to determine differences between sex in the three conditions (room temperature, 30°C, and 26°C) (D-F, H, I, K, and M). RT, room temperature; BAT, brown adipose tissue; scWAT, subcutaneous white adipose tissue; HU, Hounsfield units.

**Figure 6.**
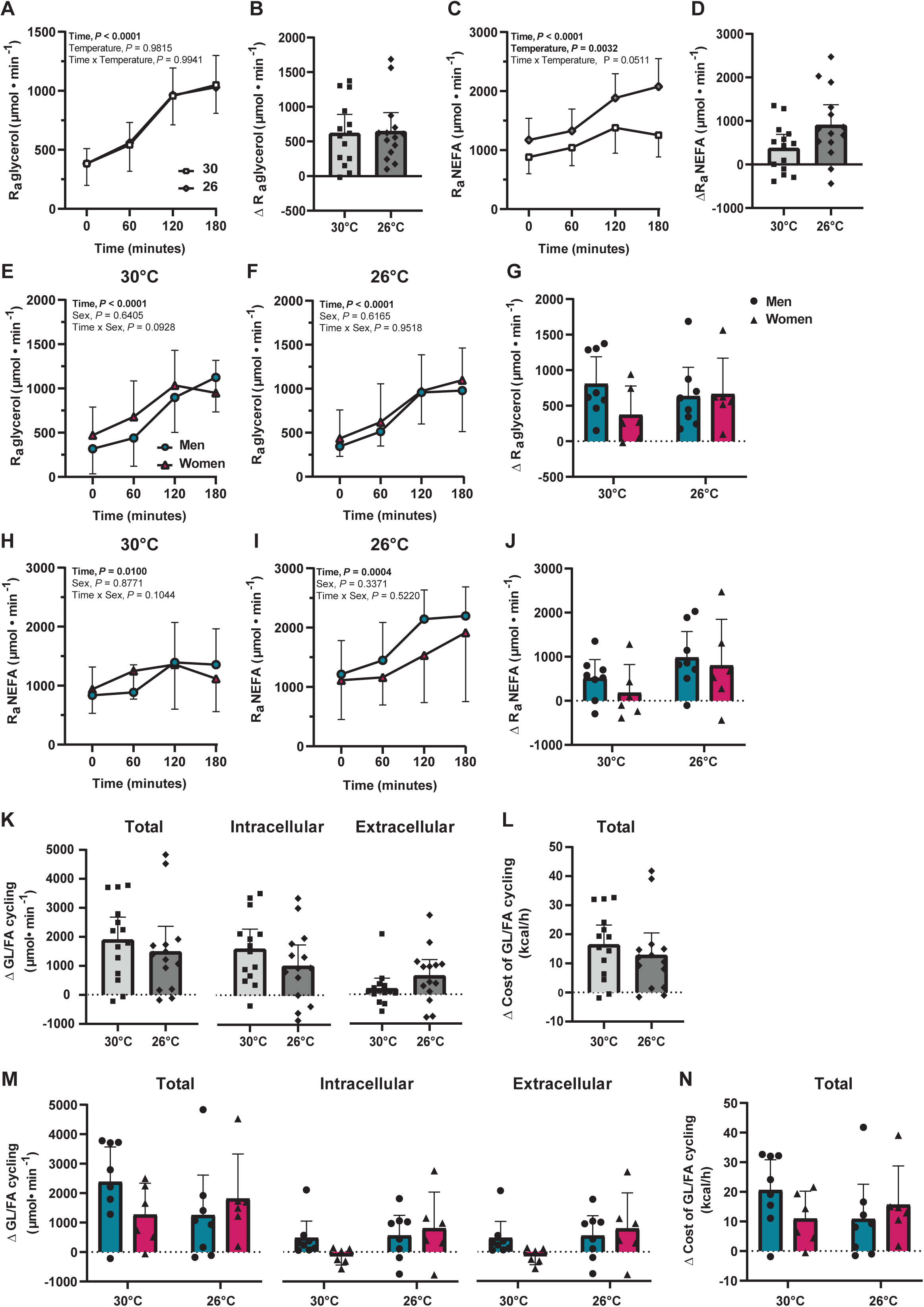
Degree of cold exposure does not modulate white adipose tissue lipolysis and glycerolipid-fatty acid cycling. (A and B) Systemic rate of appearance of glycerol over time (A) and cold-stimulated change (B) in response to skin temperature of either 30°C or 26°C among all participants (30°C: n = 14, 26°C: n = 14). (C and D) Systemic rate of appearance of NEFA over time (C) and cold-stimulated change (D) in response to skin temperature of either 30°C or 26°C among all participants (30°C: n = 14, 26°C: n = 14). (E-G) Systemic rate of appearance of glycerol over time (E-F) and cold-stimulated change (G) in response to skin temperature of either 30°C or 26°C in men (30°C: n = 8, 26°C: n = 8) and women (30°C: n = 6, 26°C: n = 6). (H-J) Systemic rate of appearance of NEFA over time (H-I) and cold-stimulated change (J) in response to skin temperature of either 30°C or 26°C in men (30°C: n = 8, 26°C: n = 8) and women (30°C: n = 6, 26°C: n = 6). (K and M) Cold-stimulated change in total, intracellular, and extracellular glycerolipid/fatty acid (GL/FA) cycling at skin temperature of either 30°C or 26°C among all participants (30°C: n = 14, 26°C: n = 14) (K), and in men (30°C: n = 14, 26°C: n = 14) and women (30°C: n = 14, 26°C: n = 14) (M). (L and N) Cold-stimulated change in the energetic cost of total GL/FA cycle in response to skin temperature of either 30°C and 26°C among all participants (30°C: n = 14, 26°C: n = 14) (L), and in men (30°C: n = 8, 26°C: n = 8) and women (30°C: n = 6, 26°C: n = 6) (N). Data are reported as means with 95% confidence interval (CI). P-values are shown only for significant differences (i.e. P < 0.05). A two-way ANOVA for repeated measures with Bonferroni post hoc test was used to determine the effects of the time, the temperature and their interaction (A, C, E, F, H, I). A mixed-model ANOVA with Bonferroni post hoc test was used to determine differences between sex. The difference between a skin temperature of 30°C and 26°C determined using paired-sample t-test (B, D, K and L).

### The cold exposure intensity does not modulate white adipose tissue lipolysis and glycerolipid- fatty acid cycling

To investigate the effects of skin cooling on WAT lipolysis and glycerolipid-fatty acid futile cycling, we used a continuous infusion of [U-^13^C]palmitate and [1,1,2,3,3-^2^H_5_]glycerol to determine the systemic rate of appearance (R_a_) of NEFA (Ra_NEFA_) and glycerol (Ra_glycerol_) (an indicator of whole-body WAT lipolysis). The systemic Ra_glycerol_ and Ra_NEFA_ both increased significantly in the cold (*P* < 0.001; Figures 6A and C), but Ra_NEFA_ increased by a greater magnitude at 26°C than at 30°C (time by condition interaction, *P* = 0.051). The rate of fatty acid oxidation mirrored the increase in the systemic Ra_NEFA_. Compared to room temperature, the rate of fatty acid oxidation increased both at a mean skin temperature of 30°C and 26°C but increased to a greater degree at 26°C (Table 2, time by condition interaction, *P* = 0.0001). In contrast, the rate of carbohydrate utilization did not change significantly at 30°C among grouped participants, whereas at mean skin temperature of 26°C, the rate of carbohydrate utilization averaged over the 60 min of the cold exposure was significantly higher compared to room temperature (989 μmol·min^-1^ [95% CI: 724 to 1253] vs. 1827 μmol·min^-1^ [95% CI: 1339 to 2316]; *P* < 0.001) and compared to the mean skin temperature of 30°C (1206 μmol·min^-1^ [95% CI: 593 to 1818]; *P* < 0.0001) (Table 2). Under steady-state conditions, the difference between the total fatty acids released upon intracellular hydrolysis of triglycerides (3× Ra_glycerol_) and the rate of total fatty acid oxidation (calculated from indirect calorimetry) indicate the whole-body rate of fatty acid re- esterification, as this is primarily its alternative metabolic fate. The results show that there was no difference in fatty acid re-esterification or the metabolic cost of cycling between the two skin cooling conditions or between sexes (Figure 6K-N). This suggests that, like BAT oxidative metabolism, glycerolipid-fatty acid cycling follows an all-or-none response and any increase in the release of fatty acids in circulation is matched by an increase in fatty acid oxidation.

### Metabolites and hormones change during cold exposure

The effects of cold exposure on changes in metabolite and hormone concentrations are presented in Table 2. C-peptide, TSH and free T_4_ levels decreased in the cold, regardless of the temperature of skin cooling (effect of temperature, P < 0.002). Temperature (room temperature *vs.* cold exposure) by condition (30°C vs. 26°C) interactions were observed in glycemia, insulinemia, NEFA concentrations, palmitate, oleate and linoleate levels, GIP, cortisol and TSH. Temperature by condition by sex interactions were observed for glucagon levels (*P* = 0.037), with levels decreasing at 30°C in men in particular (*P =* 0.025). Finally, sex-based differences were observed exclusively in NEFA, palmitate, oleate, and linoleate concentrations, as well as free T_3_ and free T_4_ levels.

## DISCUSSION

In the past decades, there has been considerable interest in understanding the thermogenic mechanisms driving cold-stimulated heat production in humans. Although much of the focus has been directed at stimulating BAT^7,35,38–41^, tonic contractions of postural muscles and overt shivering^38^ remain necessary for generating sufficient heat to counteract heat loss. These studies have also highlighted the vast inter-individual variability in whole-body heat production in response to mild cold stimulation. Some of this variability can be attributed to the wide range of experimental designs across studies, which creates variability in skin cooling thereby impacting the recruitment of cold-defense responses. To address this critical gap and investigate the relationship between shivering and NST in adipose tissue in response to different levels of skin cooling, we used a closed-loop temperature control system designed to carefully maintain the mean skin temperature at 30°C or 26°C. We also examined whether there were any effects on myocardial metabolism and any sex-related discrepancies in the contribution of these thermogenic mechanisms to whole-body heat production. As expected, whole-body heat production increased as a function of skin cooling, which was driven primarily by a temperature-dependent increase in shivering intensity and myocardial oxidative metabolism. In contrast, adipose tissue NST increased in response to skin cooling but was not influenced by the degree of cold stress as their thermogenic activities remained similar between 30°C and 26°C. Finally, sex related differences were only seen in shivering intensity and myocardial oxidative metabolism, with women shivering to a greater degree at the colder temperature mainly through the preferential recruitment of lower body muscles, and presenting with higher myocardial oxidative metabolism at all temperatures.

The two skin cooling temperatures were selected to elicit a cold stress sufficient to stimulate shivering while limiting movement to allow for the acquisition of PET images. By maintaining a mean skin temperature of either 30°C or 26°C, energy expenditure increased by 26% (95% CI: 12 to 41 %) and 100% (95% CI: 83 to 117 %), respectively; both considered a mild cold stimulus^42^. Although energy expenditure for the same mean skin temperature was greater in men compared to women, this difference disappeared when adjusting for sex-related differences in body mass. Further, the cold-induced increase in thermogenesis was also the same between sexes, which is consistent with previous observations using similarly mild conditions whether clamping skin temperature^6^, using a fixed water temperature of 18°C circulating through a cooling suit^35^, through investigator-determined subjective cooling^7,43^, or using a fixed air temperature^44^. It has long been known that tonic muscle contractions and overt shivering drives this cold-stimulated increase in heat production, but it has often been postulated that this may be less so under mild cold conditions^45,46^. Here we show that even under mild conditions, shivering intensity increases as a function of skin cooling, with women shivering to a greater extent than men^43,47^. This sex- related difference was largely explained by differences in lean body mass, which suggests that women needed to recruit a higher proportion of muscle fibers to produce the same amount of heat. Interestingly, for a same skin cooling temperature, there was tremendous inter-individual variability in the shivering intensity (95% CI: 0.9 to 2.4 %MVC at T_skin_ of 30°C; 95% CI: 3.0 to 5.7 %MVC at T_skin_ of 26°C). This variability may be explained by the various shivering patterns exhibited between individuals, including differences in the recruitment of certain muscle groups or specific subpopulations of fibers within the same muscle (distinguished by burst frequency and intensity)^38,48,49^.

The characterization of shivering under cold stimulation is often limited to the magnitude of muscle contractions (*e.g.* percentage of maximal voluntary contractions), but very little is known about the shivering pattern (muscle recruitment, continuous vs burst shivering, burst frequency). This is particularly relevant since the shivering pattern can significantly impact metabolic fuel selection (carbohydrate vs. lipid)^3,50,51^, motor control^52,53^ and thermal comfort. Using surface EMG, we observed a preferential recruitment of large centrally located muscles like *m. trapezius, m. pectoralis major, m. deltoideus, m. sternocleidomastoid* at 30°C but a progressive reliance on larger lower body muscles like the quadriceps at 26°C. This was most evident in women, who presented a higher shivering intensity in the three major muscles of the quadriceps. The frequency and intensity of burst shivering increased as a function of skin cooling, as did the intensity of continuous shivering. However, only the burst shivering intensity differed between sexes, and only when skin temperature was cooled to 26°C (14.4 %MVC for women vs 8.7 %MVC for men). This increase in burst intensity explains the greater overall shivering intensity in women, which can impact fine motor control^52^ and may result in increased discomfort in the cold, despite women reporting similar thermal sensation to men. Indeed, shivering intensity, burst frequency, burst intensity and continuous intensity were all associated with thermal sensation, which was mainly driven by a strong association in women (r = -0.90, *P <* 0.0001; r = -0.94, *P* < 0.0001; r = -0.83, *P* = 0.0008, r = -0.91, *P* < 0.0001, respectively).

While significant gaps remain in our understanding of the induction and control of shivering and its impact on cold-induced thermogenesis, even less is known about the range between basal and maximal activity of non-shivering mechanisms in either animals or humans. The classic study by Foster and Frydman remains to date the only preclinical study to provide indirect estimates of the relative contribution of BAT thermogenesis on whole-body heat production at different temperatures (-19°C, -6°C, 6°C, and at thermoneutrality)^54^. They showed that, in warm-acclimated rats – a model that more closely resembles human conditions – both the absolute blood flow and the percentage of cardiac output directed towards BAT increases in response to the cold (6°C) but does not increase further with decreasing ambient temperatures. Similarly, interscapular BAT temperature of Siberian hamsters does not differ whether exposed to ambient temperature (21°C) or to a more severe cold stress (4°C)^55^. Here, we provide the first direct measures of BAT oxidative metabolism in response to two different skin cooling conditions. We show that BAT oxidative metabolism and glucose uptake increase in response to mild cold exposure but does not increase further in response to a colder stimulus. This is consistent with studies using [^15^O]O_2_ PET to quantify BAT oxygen consumption, which reported similar rates of oxygen consumption in healthy individuals regardless of the cold stimulus^20,30^, so long as the cold stress meets a minimal threshold. Similarly, Ra_glycerol_, an indicator of whole-body adipose tissue lipolysis, increased in response to the cold but did not differ between skin cooling conditions. Although the systemic Ra_NEFA_ increased as a function of skin temperature, it was matched by an increase in the rate of fatty acid oxidation. The net result was a plateauing of another adipose tissue derived form of thermogenesis, the recruitment of the GL/FA cycling. Evidence suggests that there may be selective control of the sympathetic outflow, which can result in target-specific differential responses to thermal stimulations^56–58^. The preclinical findings and our current study suggest that with a minimal cold stimulus to trigger the sympathetic nervous system, the release of norepinephrine may be sufficiently high to saturate the adrenergic receptors on adipose tissue. This would result in WAT lipolysis and BAT thermogenesis being readily maximally stimulated rather than responding in a graded fashion. This type of response can also be observed in cold-stimulated sympathetic nervous system mediated vasoconstriction, where vascular conductance and skin sympathetic nerve activity seem to reach a plateau of activity in response to the cold ^59,60^. We therefore contend that under cold stimulation, the major determinant of BAT thermogenesis in an individual is the tissue’s recruitment state at the time of measurement (*e.g.* atrophied/*whitened* from de-acclimation *vs*. increased recruitment from intermittent or chronic cold exposure)^61–63^ or the number of sympathetic ganglion cells innervating BAT which impacts the amplitude of the BAT sympathetic response^64^. It does not appear to be determined by inter-individual variations in the degree of activation (*i.e.* the tissue is either fully activated or basally active). This does not imply that, within the same individual, BAT cannot be activated to different degrees, as it may indeed differ in thermogenic activity in a stimulus-dependent fashion. For instance, in the same individual, cold-stimulated BAT oxidative metabolism is nearly 6-times higher than when pharmacologically stimulated using a single maximum allowable dose of mirabegron (200 mg)^25^. This suggests that mirabegron at this dose may not fully saturate the adrenergic receptors on the surface of brown adipocytes like norepinephrine might under cold stimulations.

Beyond the thermogenic responses to the cold, we were equally interested in examining whether cardiovascular responses to cold exposure followed a dose-response. Systolic and diastolic blood pressure increased in a similar manner (∼10 mmHg) for both mean skin temperatures, consistent with the SNS-mediated vasoconstriction^65–67^. Heart rate decreased by 4 beats·min^-1^ at the mean skin temperature of 30°C, as expected. However, at the mean skin temperature of 26°C, it surprisingly increased by 3 beats·min^-1^. Cold exposure often, though not always^68^, results in a baroreceptor reflex-mediated decrease in heart rate in both younger and older individuals yet the observed increase in our study is not commonly reported^65–67^. The increased heart rate was also accompanied by an increase in rate pressure product, a sensitive index of myocardial oxygen consumption. This was consistent with the increase in myocardial oxidative metabolism during the coldest skin cooling condition, measured using [^11^C]-acetate PET. Interestingly, both shivering thermogenesis and myocardial oxidative metabolism increased progressively in the cold and show a moderate association (r = 0.45, *P* = 0.02), whereas BAT oxidative metabolism was not associated with myocardial oxidative metabolism (r = 0.18, *P* = 0.36). Myocardial [^11^C]-acetate uptake rate was also significantly higher at the coldest skin temperature, which could be explained by a higher myocardial perfusion^69^. Interestingly, all three myocardial parameters were significantly higher in women than men. Collectively, these cardiometabolic results suggest that shivering and myocardial thermogenesis may be associated but also that the sympathetic efferents that regulate adipose tissue lipolysis and thermogenesis are likely distinct from cardiovascular sympathetic efferents or the latter may be modulated by baroreceptor control of blood pressure.

## LIMITATIONS OF THE STUDY

Although we provide novel insights into the integrated response between shivering and NST in adipose tissue under two levels of skin cooling conditions in both men and women, this study also has certain limitations. First, the relatively small sample size, combined with the variability in certain outcome measures limit the statistical power to be more affirmative about certain findings beyond the primary outcomes of adipose tissue NST. Second, we included unacclimatized young, healthy men and women, which limits our ability to determine whether the ST and NST in adipose tissue are modulated following acclimation/acclimatization, in individuals with a higher percentage of adiposity or with metabolic disorder, across ethnicities or in an older cohort. Third, although we clamped the skin temperature to minimize inter-individual variation in cold responses, we still observed variability in heat production. This variability may be due to differences in the sensitivity and/or density of cold-sensitive cutaneous thermoreceptors, variations in the integration of visceral thermosensory feedback, or differences in body morphology and composition among other factors.

## CONCLUSION

In summary, we simultaneously examined shivering and adipose tissue NST at two different mean skin temperatures. By standardizing the cooling stimulus using a novel mean skin temperature clamping technique, this study demonstrates that cold-stimulated heat production is driven to a large extent by shivering and myocardial thermogenesis in men and women. The increase in myocardial oxidative metabolism appears to be dependent upon the intensity of skin cooling and was associated with the increase in shivering intensity. Conversely, adipose tissue NST does not respond linearly to skin cooling, in contrast to common perception. Finally, sex related differences were only seen in the cold-induced changes in heat production and shivering intensity, with women generating less heat than men, yet shivering to a greater degree at colder temperatures primarily through their lower body muscles and through a greater burst shivering intensity. Myocardial oxidative metabolism and therefore its thermogenic contribution in the cold was also greater in women than men, at all temperatures. Further work is needed to determine the extent to which skeletal muscle NST or other forms of futile cycles impact the variability in cold- induced heat production or shivering patterns in men and women.

## RESOURCE AVAILABILITY

### Lead contact

Further information and requests for resources and reagent should be directed to and will be fulfilled by the lead contact, Denis P. Blondin.

### Data and code availability

The processed data generated during this study can be made available upon request to the lead contact following internal review and established data protection laws. The MATLAB code used to perform the pharmacokinetic modeling of FDG and [^11^C]-acetate are available at doi.org/10.5281/zenodo.5834789..

## ACKNOWLEDGEMENTS

The authors would like to thank the participants of this study for their commitment and collaboration. The authors thank Caroll-Lynn Thibodeau, Maude Gérard, Myriam Flipot, Jennefer- Ann Lefrançois, Éric Lavallée, Esteban Espinosa, Christophe Noll and Lucie Bouffard for their excellent technical assistance and Stephen C. Cunnane for the use of PMOD. D.P. Blondin holds the GSK Chair in Diabetes of the Université de Sherbrooke and a Fonds de Recherche du Québec- Santé (FRQS) J2 salary award. L. Dumont is the recipient of an FRQS doctoral training award. A.C. Carpentier is supported by a Canada Research Chair in the Molecular Imaging of Diabetes. This work was supported by a Natural Sciences and Engineering Research Council of Canada Discovery Grant (RGPIN-2019-05813) and Discovery Launch Supplement (DGECR-2019- 00049), an FRQS Starting Grant for New Investigators – Junior 1 and an equipment grant from the Canada Foundation for Innovation John R. Evans Leaders Fund (JEFL-43148) to D.P. Blondin.

## AUTHORS CONTRIBUTION

Conceptualization, D.P.B; Methodology, L.D., D.P.B, R.E., A.S., R.F., J.B. and A.C.C.; Investigation, L.D., G.R., E.C., M.F., F.F., S.P., S.D., B.G., E.E.T., A.C.C., and D.P.B.; Writing first draft, L.D. and D.P.B; Writing – review and editing, L.D., G.R., E.C., M.F., F.F., S.P., S.D., B.G., E.E.T., R.E., R.F., A.S., J.B., A.C.C., and D.P.B.; Visualization, L.D. and D.P.B; Funding Acquisition, D.P.B.

## STAR METHODS

### KEY RESOURCES TABLE

2 KEY RESOURCES TABLE
| REAGENT or RESOURCE | SOURCE | IDENTIFIER |
| --- | --- | --- |
| **Biological Samples** |  |  |
| Human plasma samples | This paper | N/A |
| Human urine samples | This paper | N/A |
| **Chemicals, Peptides, and Recombinant Proteins** |  |  |
| C-peptide, GIP, GLP-1 total,<br>Glucagon, Insulin, Leptin | Sigma-Millipore | HMH3-34K-07 |
| Aprotinin | Sigma-Aldrich | A6279 |
| **Critical commercial Assays** |  |  |
| Glycerol reagent set | Instruchemie, Sigma-Aldrich | Cat# 2913, Cat# F6428 |
| NEFA C kit | Wako diagnostics, Randox<br>laboratories | Cat# 434-91795, Cat# FA115 |
| **Software and Algorithms** |  |  |
| GraphPad Prism 8 for statistical<br>analysis | GraphPad Software | RRID: SCR_002798 |
| Adobe Photoshop | Adobe | https://www.adobe.com/ |
| PMOD 3.7 | PMOD Technologies LLC | RRID : SCR_016547 |
| MATLAB R2019a | The MathWorks | RRID : SCR_001622 |
| Python | Python Software Foundation | N/A |
| Grafana | Grafana Labs | N/A |
| **Other** |  |  |
| [<sup>11</sup>C]acetate, radiotracer | CIMS | N/A |
| [<sup>18</sup>F]FDG, radiotracer | CIMS | N/A |
| [3-<sup>3</sup>H]glucose, stable isotope<br>tracer | PerkinElmer | Cat# NET331A005MC |
| [U-<sup>13</sup>C]palmitate, stable isotope tracer | Cambridge isotopes | Cat# CLM-3943-MPT |
| [1,1,2,3,3-<sup>2</sup>H<sub>5</sub>]glycerol, stable isotope tracer | Cambridge isotopes | Cat# DLM-1229-MPT |
| Luminex xMAP-based immunoassays | Millipore | N/A |
| GC-MS/MS | Waters | XevoTQ-GC |
| Myomonitor IV EMG system | Delsys Inc | N/A |
| eCelsius® Performance electronic capsule | BodyCAP | https://www.bodycap.us/ |
| ePerf® Connect Gateway & ePerf Mobile application | BodyCAP | https://www.bodycap.us/ |
| Liquid-conditioned suit | Med-Eng Inc. | Cat# 1004340 |
| Isotemp 6200R28 water bath | Fisher Scientific | Cat# 13-874-676 |
| FlowKit-500, mass flow controller | Sable System International | N/A |
| FMS Field Metabolic System, Respirometry system | Sable System International | N/A |
| Horizon QDR DXA | Hologic | N/A |
| Biograph Vision 600 PET-CT scanner | Siemens Healthineers | N/A |
| Raspberry Pi 3B+ | Raspberry Pi | N/A |
| Temperature sensors | TE Connectivity | Cat# NB-PTCO-153 |

### EXPIRIMENTAL MODEL AND PARTICIPANT DETAILS

#### Human Participants

Eight healthy adult men with a mean age of 28 years (95% CI: 24-31), body mass index of 25.3 kg/m^2^ (95% CI: 23.6-26.9) and body surface area of 2.08 m^2^ (95% CI: 1.99-2.17) and six healthy adult women with a mean age of 26 years (95% CI: 20-32), body mass index of 21.9 kg/m^2^ (95% CI: 19.2-24.5) and body surface area of 1.65 m^2^ (95% CI: 1.50-1.80) volunteered to participate in two metabolic experimental sessions (Table 1). We have documented the contraceptive methods used by the women under study, despite the fact that the menstrual cycle has no effect on thermogenic response^70^. All participants were informed of the methodology, and informed written consent was obtained in accordance with the Declaration of Helsinki. The protocol was approved by the Human Ethics Committee of the Centre de recherche du Centre hospitalier universitaire de Sherbrooke (protocol #: 2021-3978). The inclusion criteria included: (i) age between 18 and 35 years; (ii) BMI < 29.9 kg/m^2^; (iii) normal fasting glucose (< 5.6 mmol/l); (iv) normal glucose tolerance (2 h post 75 g OGTT glucose < 7.8 mmol/l); (v) HbA1c < 5.8%. Exclusion criteria included: (i) overt cardiovascular disease as assessed by medical history, physical exam, and abnormal ECG; (ii) treatment with any drug known to affect lipid or carbohydrate metabolism; (iii) presence of liver disease, uncontrolled thyroid disorder, previous pancreatitis, bleeding disorder, or other major illness; (iv) smoking (>1 cigarette/day) and/or consumption of 2 alcoholic beverages per day; (v) prior history or current fasting plasma cholesterol level > 7 mmol/l or fasting TG > 6 mmol/l; (vi) women who are pregnant or breastfeeding.

**Table 1.**
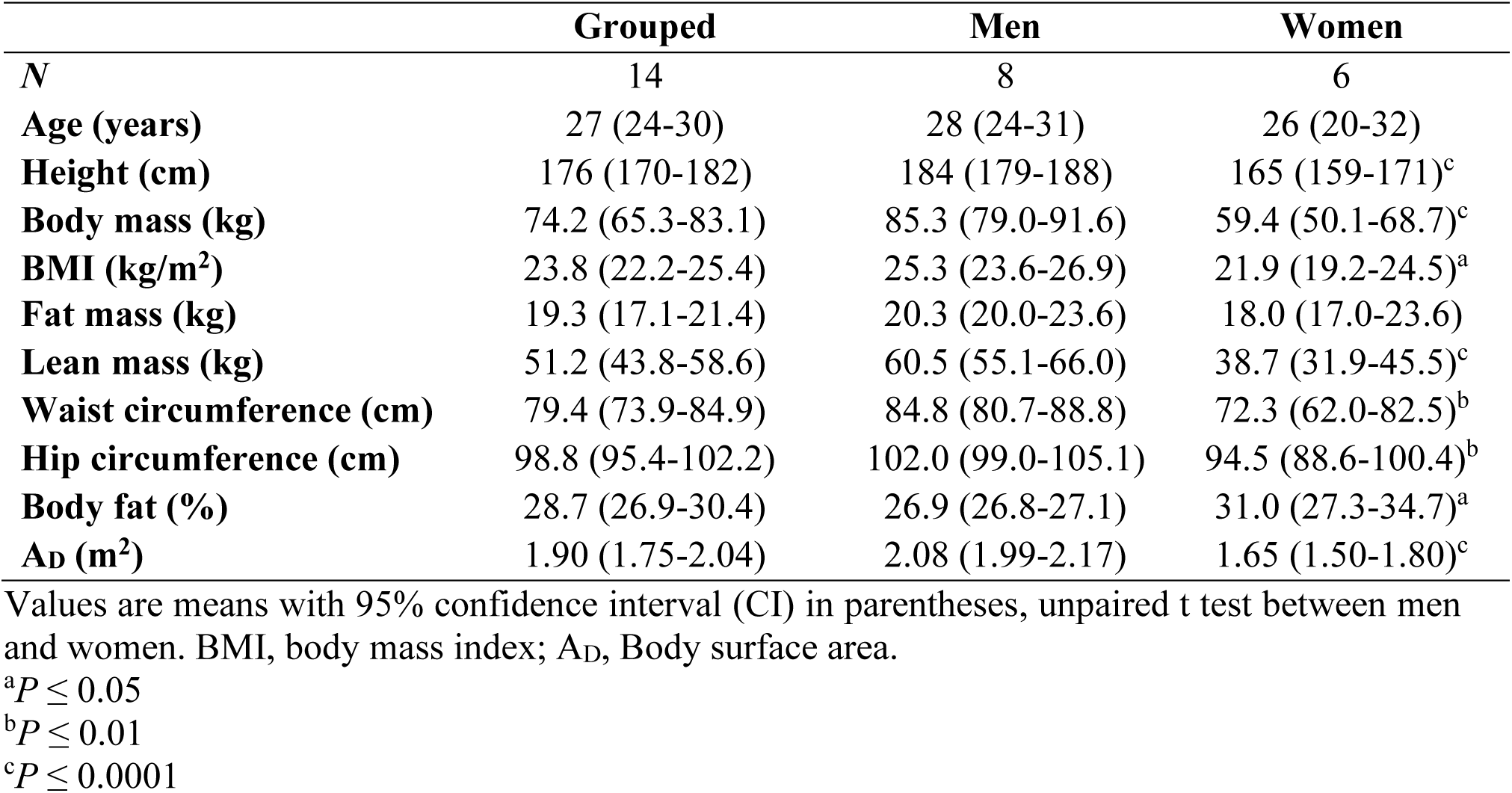
Participant characteristics.

|  | **Grouped** | **Men** | **Women** |
| --- | --- | --- | --- |
| *N* | 14 | 8 | 6 |
| **Age (years)** | 27 (24-30) | 28 (24-31) | 26 (20-32) |
| **Height (cm)** | 176 (170-182) | 184 (179-188) | 165 (159-171)<sup>c</sup> |
| **Body mass (kg)** | 74.2 (65.3-83.1) | 85.3 (79.0-91.6) | 59.4 (50.1-68.7)<sup>c</sup> |
| **BMI (kg/m<sup>2</sup>)** | 23.8 (22.2-25.4) | 25.3 (23.6-26.9) | 21.9 (19.2-24.5)<sup>a</sup> |
| **Fat mass (kg)** | 19.3 (17.1-21.4) | 20.3 (20.0-23.6) | 18.0 (17.0-23.6) |
| **Lean mass (kg)** | 51.2 (43.8-58.6) | 60.5 (55.1-66.0) | 38.7 (31.9-45.5)<sup>c</sup> |
| **Waist circumference (cm)** | 79.4 (73.9-84.9) | 84.8 (80.7-88.8) | 72.3 (62.0-82.5)<sup>b</sup> |
| **Hip circumference (cm)** | 98.8 (95.4-102.2) | 102.0 (99.0-105.1) | 94.5 (88.6-100.4)<sup>b</sup> |
| **Body fat (%)** | 28.7 (26.9-30.4) | 26.9 (26.8-27.1) | 31.0 (27.3-34.7)<sup>a</sup> |
| **Ad (m<sup>2</sup>)** | 1.90 (1.75-2.04) | 2.08 (1.99-2.17) | 1.65 (1.50-1.80)<sup>c</sup> |

Values are means with 95% confidence interval (CI) in parentheses, unpaired t test between men and women. BMI, body mass index; Ad, Body surface area.
<sup>a</sup> $P \leq 0.05$
<sup>b</sup> $P \leq 0.01$
<sup>c</sup> $P \leq 0.0001$

**Table 2.**
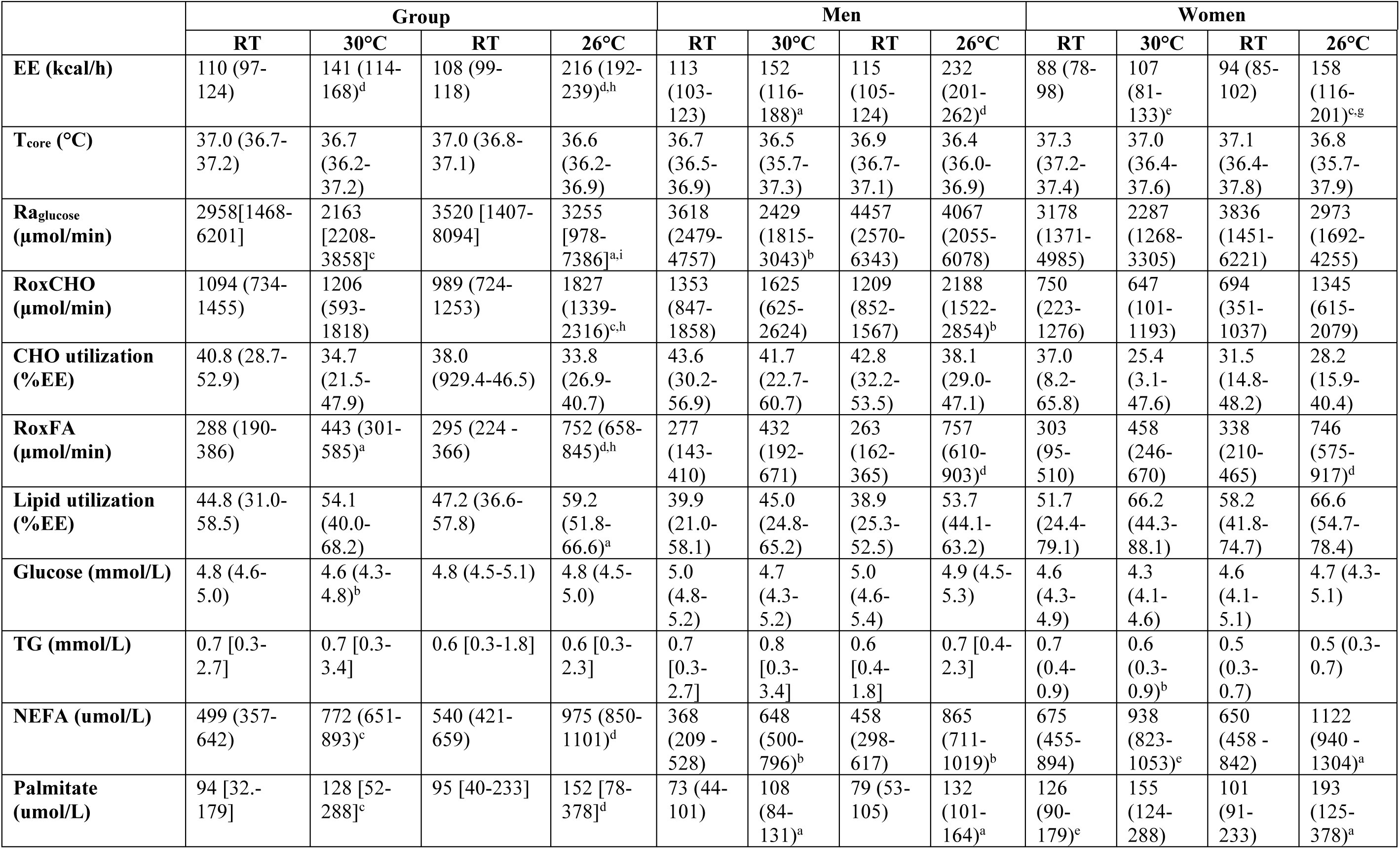
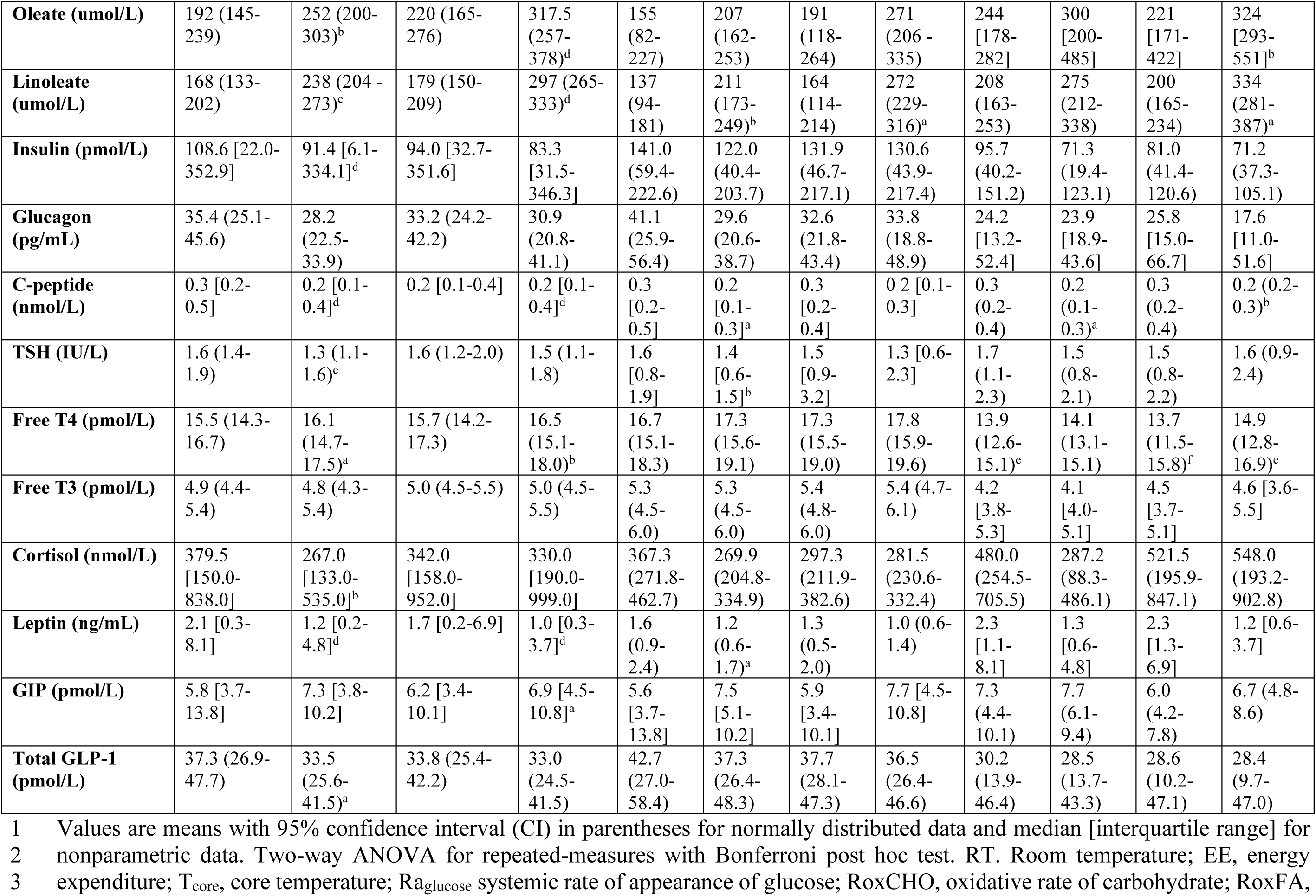
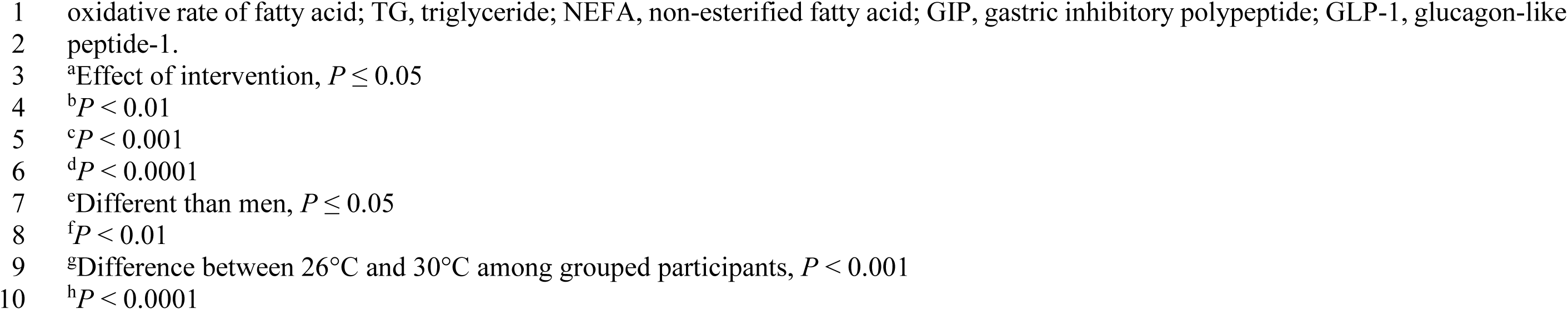
Whole-body response to cold exposure at room temperature and averaged over final 60 min of cold exposure.

### METHOD DETAILS

#### Experimental Procedures

Participants took part in two experimental sessions of acute cold exposure assigned in a random order (Figure 1). Each experimental session consisted of a 150 min baseline period at ambient temperature (∼22°C), followed by 180 min of cold exposure, elicited using a liquid conditioned suit (LCS; Three Piece, Med-Eng) perfused with cold water to reach two different mean skin temperatures: 30°C and 26 °C. We used a custom-designed feedback control loop with a temperature and flow-controlled circulation bath (Isotemp 6200R28; Fisher Scientific Co., Pittsburgh, PA, USA) to circulate water at a fixed rate and quickly elicit the stable mean skin temperature of either 30°C or 26°C. The water circulating through the LCS was rapidly adjusted and the water temperatures ranging from -5.5°C to 26.8°C (see description of skin cooling control system below). The same suit was used for all subjects to maintain consistent tubing density and water flow.

Experimental sessions were conducted between 07.00 and 15.00 h at the Centre de recherche du Centre hospitalier universitaire de Sherbrooke, following a 12 h fast and 48 h without strenuous physical activity. Participants followed a 2-day standard isocaloric diet that was determined following 3-day food record, accounting for the participant’s standard daily physical activity level, determined by wearing an accelerometer device (ActiGraph; GT3X) on top of the dominant (right or left) hip for 3-days. Upon arriving to the laboratory, participants emptied their bladder, then participant only wearing shorts and sports bras for women were weighed. Participants were instrumented with 12 temperature sensors and 8 surface electromyography electrodes (Myomonitor IV, Delsys Inc, Natick, USA) and perform a series of muscle contractions (MVC) of each of the muscles being measured for shivering activity. Participants were then fitted with the LCS, swallowed a telemetric thermometry capsule to measure core temperature and indwelling catheters were placed in the antecubital vein in both arms for blood sampling and tracer infusion.

#### Computer-controlled closed-loop skin temperature control system

To stabilize and maintain the mean skin temperature at a predetermined value, we developed a computer-controlled closed-loop skin temperature control system. A water bath was initially set to a temperature of -10°C to rapidly reach our target temperature and a 50:50 glycol-water mixture was circulated through the suit to begin the cooling. Skin temperature was measured every three seconds using twelve custom-designed temperature sensors with a sensitivity of 0.2°C. The sensors were purposefully fabricated and insulated to minimize bias caused by the temperature of the circulating water, thereby providing accurate skin temperature measurements. Mean skin temperature was monitored continuously using the twelve temperature sensors fixed to the skin, calculated using an area-weighted equation from 12 sites^72^: forehead (7%), chest (9.5%), biceps (9%), forearm (7%), abdomen (9.5%), lower back (9.5%), upper back (9.5), front calf (8.5%), back calf (7.5%), quadriceps (9.5%), hamstrings (9.5%), and hand (4%), The temperature of each site was sent to a single-board computer (Raspberry Pi) and then the mean skin temperature was calculated and displayed using an interactive visualization web application (Grafana). A custom algorithm programmed in Python used a proportional-integral-derivative (PID) controller (PID parameters: Kp = 1, Ki = 0.1, Kd = 0.05, T_min_ = -10°C and T_max_ = 50°C) to control the temperature of the circulating glycol-water mixture to reach the target mean skin temperature. The setpoint was updated autonomously every 30 seconds and a command was sent to the bath water system to maintain a set temperature. The temperature measured by each sensor and the area-weighted mean skin temperature collected every 3 seconds were sent to a database for real-time monitoring and subsequent analysis.

#### Muscle recruitment

Shivering EMG signals were recorded from 8 muscles *m. trapezius superior* (TS), *m. sternocleidomastoid* (SCM), *m. pectoralis major* (PM), *m. deltoideus* (DT), *m. vastus lateralis* (VL), *m. rectus femoris* (RF), *m. vastus medialis* (VM) and *m. bicep femoris* (BF). Surface electrodes were placed over the bellies of each muscle. Raw EMG signals were collected at 1000 Hz, filtered to remove spectral components below 20 Hz and above 500 Hz as well as 60 Hz contamination and related harmonics, and analyzed using custom designed algorithms. Shivering activity of the 8 individual muscles was monitored 10 min before and continuously throughout cold exposure. Shivering intensity of individual muscle was determined from root-men-square (RMS) values calculated from raw EMG, as previously described ^49,71^. Voluntary muscle activity was minimized throughout cold exposure by asking participants to avoid voluntary movements during the recording periods.

#### Thermal responses

Core temperature (Tcore) was recorded using an ingested telemetric pill which measures the temperature on the intestine each 30 sec (eCelsius Performance electronic capsule; BodyCap). Mean skin temperature (Tskin) was monitored continuously using 12 temperature sensors fixed to the skin, calculated using an area-weighted equation from 12 sites: forehead, chest, biceps, forearm, abdomen, lower and upper back, front and back calf, quadriceps, hamstrings and hand^72^. Subjective thermal sensation was determined every 30 min by asking participants about their perception of their temperature using a 11-point Likert scale ranging from -5 (“coldest ever experienced”) to +5 (“warmest ever experienced”), and 0 being defined feeling neither cold or warm. Thermal responses, shivering activity, metabolic rate and substrate utilization were measured continuously during the final 30 min of baseline and the subsequent 180 min of cold exposure.

#### Metabolic Heat Production and Fuel Selection

Whole-body metabolic heat production and substrate utilization was determined by indirect respiratory calorimetry (corrected for protein oxidation), measured continuously 30 min before and for the entire duration of cold exposure. The molar rate of fatty acid oxidation (µmol·min^-1^) was calculated from triglyceride oxidation (g·min^-1^) assuming the average molecular weight of triglyceride as 861 g·mol^-1^ and multiplying the molar rate of triglyceride oxidation by three, as three moles of fatty acids are contained in each mole of triglyceride. Energy potentials of 16.3, 40.8 and 19.7 kJ·g^-1^ were used to calculate the amount of heat produced from glucose, lipid and protein oxidation, respectively.

#### PET Imaging and Analysis

Participants remained supine in a PET and CT scanner (Siemens Biograph Vision 600) for 30 min for the baseline and the final 90 min of the cold exposure. Tissue-specific oxidative metabolism was determined by first performing a CT scan (30 mAs) centered at the cervico-thoracic region to correct for attenuation and define PET regions of interest (ROI). At time -30 min (room temperature, before cold exposure) and at time 90 min, a ∼ 175 MBq bolus of [^11^C]-acetate was injected intravenously and was followed by a 20-min list-mode dynamic PET acquisition (frames: 24 x 10 s, 13 x 30 s, 2 x 300 s) centered at the cervico-thoracic junction^39^. A ∼125 MBq bolus of FDG was intravenously injected 30 min after the [^11^C]-acetate injection at time 120 followed by a list-mode dynamic PET acquisition (frames: 12 x 10 s, 10 x 45 s, 7 x 90 s, 4 x 300 s) centered at the cervico-thoracic junction. Residual [^11^C]-acetate activity prior to the FDG scans were corrected by acquiring a 30 s frame prior to the injection of FDG and accounting for the disintegration rate and metabolic clearance of ^11^C.

ROI were drawn from the transaxial CT slices then copied to the [^11^C]-acetate and FDG PET images. ROIs were drawn in the left ventricle for blood activity (image-derived arterial input function, AIF), the larger skeletal muscles in the field of view (e.g. *m*. *sternocleidomastoid, m. trapezius, m. pectoralis major, m. deltoideus, m. levator scapulae, m. latissimus dorsi, m. erector spinea*), on posterior cervical subcutaneous adipose tissue, on supraclavicular BAT and on the left ventricle of the myocardium. The mean standard uptake values (SUV_mean_) from these ROIs were then extracted for each time frame using PMOD (version 3.7, PMOD Technologies LLC) to create time-activity curves. The blood signal for [^11^C]-acetate was then corrected to exclude the contribution of metabolites^73^. The metabolite-corrected time-activity curves were then used to perform pharmacokinetic modeling in MATLAB (The mathWorks, R2019a). A four-compartment, two-tissue, model was applied to the [^11^C]-acetate signal to derive the rates of uptake (*K*_1_ in ml‧g^-1^‧min^-1^), oxidation metabolism (*k*_2_ in min^-1^) and lipid synthesis (*k*_3_ in min^-1^)^36^. Plasma and tissue time-radioactivity curves for FDG were analyzed graphically using the Patlak linearization method^39,74^ .The glucose fractional uptake (*K_i_* in min^-1^) is equal to the slope of the plot in the graphical analysis. Net glucose uptake (K_m_ in nmol‧g^-1^‧min^-1^ = (*K*_i_‧circulating glucose)‧(LC^-1^)‧(tissue density‧1000)^-1^) was obtained by multiplying *Ki* by circulating plasma glucose levels at the time of the PET image acquisition, which assumes a lump constant (LC) value of 1.14 and 1.16 compared with endogenous plasma glucose for the BAT and skeletal muscles, respectively, and corrected for a tissue density of 0.925 g/mL and 1.0597 g/mL^75,76^.

#### Whole-Body Lipolysis and Glycerolipid/Fatty Acid Cycling

Upon arriving to the laboratory, participants emptied their bladder, and indwelling catheters were placed in the antecubital vein in both arms for blood sampling and tracer infusions. A primed (3.3 x 10^6^ d.p.m.‧min^-1^) continuous infusion (0.33 x 10^6^ d.p.m.·min^-1^) of [3-^3^H]glucose was started 150 min before the start of cold exposure (time - 150 min) to determine the systemic rate of appearance of plasma glucose (Ra_glucose_)^39^. Plasma NEFA appearance rate (Ra_NEFA_) and plasma glycerol appearance rate (Ra_glycerol_) were determined with a continuous infusion of [U-^13^C]palmitate (0.01 µmol/kg/min in 100 mL 25% human serum albumin) and a primed (1.6 µmol·kg^-1^) continuous infusion (0.05 µmol/kg/min) of [1,1,2,3,3-^2^H_5_]glycerol started 60 min before the start of the cold exposure (time - 60 min)^39^. Ra_NEFA_ was calculated by multiplying the plasma palmitate appearance rate by the fractional contribution of palmitate to total plasma NEFA concentrations.

Total, intracellular and extracellular glycerolipid/fatty acid (GL/FA) cycling was calculated using Ra_glycerol_, Ra_NEFA_ and the rate of fatty acid oxidation. In brief, total GL/FA cycling was calculated as the difference between the rate of fatty acid oxidation and the total amount of fatty acids made available by the hydrolysis of intracellular triglycerides (3 x Ra_glycerol_), which is derived primarily from WAT under fasted conditions. Since glycerol kinase has a low activity in WAT, relative to other tissues, the glycerol produced following the complete hydrolysis of a triglyceride in WAT is rapidly excreted into circulation where it can serve as gluconeogenic substrate. Intracellular or primary GL/FA cycling refers to the re-esterification of fatty acids within the cell where it was hydrolyzed. This is calculated from the difference between the systemic rate of appearance of NEFA (i.e. Ra_NEFA_) and the total amount of fatty acids released following intracellular hydrolysis of triglycerides (3 x Ra_glycerol_). Extracellular or secondary GL/FA cycling refers to the re-esterification of fatty acids after it has been released into the circulation, primarily packaged by the liver into triglyceride-rich lipoproteins (very-low density lipoproteins; VLDL) or intracellular lipid droplets. This is calculated from the difference between the total rate of fatty acids oxidation and the systemic appearance of fatty acids (i.e. Ra_NEFA_). The energy cost of GL/FA cycling was calculated assuming that the re-esterification of each triglyceride requires ∼8 ATP (or ∼144 kcal·mol^-1^ of triglyceride recycled), for the activation of each of the three fatty acids (2 ATP per fatty acid), and glycerogenesis (∼2 ATP) providing the three-carbon backbone for triglyceride synthesis^77^.

#### Biological Assays

Glucose, total NEFA, TG, cortisol, TSH, free T3 and free T4 were measured using specific radioimmunoassay and colorimetric assays^39^. Plasma C-peptide, GIP, total GLP-1, glucagon, insulin, leptin and PYY were measured using Luminex xMAP-based immunoassays (Milllipore, Etobicoke, ON, Canada). Adiponectin, total and acylated ghrelin were measured by ELISA (Alpco, Salem, NH, USA). Individual plasma NEFA (palmitate, linoleate, oleate), [U- ^13^C]palmitate enrichment, [1,1,2,3,3-^2^H_5_]glycerol enrichment and [3-^3^H]glucose enrichment were measured by gas chromatography-mass spectrometry^39^.

### QUANTIFICATION AND STATISTICAL ANALYSIS

#### Statistical Analyses

Statistical analysis was performed using Prism (GraphPad; 8.0) and Jamovi (v. 2.3.21). Blood chemistry data are expressed as mean with 95% CI or median with interquartile range, whereas data in figures are expressed as mean with 95% CI. The Shapiro-Wilk test was performed to verify the normality of data, when necessary. Two-way repeated-measures ANOVA was used to examine the main effects of temperature (room temperature, cold exposure), main effects of condition (30°C, 26°C) and their interaction whereas a three-way mixed model ANOVA was used to include the additional effects of sex (men, women). Bonferroni post-hoc test was used to examine differences in main effects and correct for multiplicity. An ANCOVA was used to correct for sex- related differences in body mass on energy expenditure. The significance threshold was set at *P* ≤ 0.05.

